# Macrophage depletion impairs neonatal tendon regeneration

**DOI:** 10.1101/2021.01.07.425735

**Authors:** Kristen L. Howell, Deepak A. Kaji, Angela Montero, Kenji Yeoh, Philip Nasser, Alice H. Huang

## Abstract

Tendons are dense connective tissues that transmit muscle forces to the skeleton. After adult injury, healing potential is generally poor and dominated by scar formation. Although the immune response is a key feature of healing, the specific immune cells and signals that drive tendon healing have not been fully defined. In particular, the immune regulators underlying tendon regeneration are almost completely undetermined due to a paucity of tendon regeneration models. Using a mouse model of neonatal tendon regeneration, we screened for immune-related markers and identified upregulation of several genes associated with inflammation, macrophage chemotaxis, and TGFβ signaling after injury. Depletion of macrophages using AP20187 treatment of MaFIA mice resulted in impaired functional healing, reduced cell proliferation, reduced ScxGFP+ neo-tendon formation, and altered tendon gene expression. Collectively, these results show that inflammation is a key component of neonatal tendon regeneration and demonstrate a requirement for macrophages in effective functional healing.

## INTRODUCTION

Tendons are dense, connective tissues that connect muscle to the skeleton (1). Injury to tendons is extremely common and results in pain and sustained loss of mechanical function (2–5). This is largely due to the poor healing capacity of tendons and inability restore native matrix architecture since tendons heal by disorganized scar formation (5, 6). Currently, treatment options remain few and the gold standard of treatment is surgical repair, which does not regenerate tendon structure or function. Identifying novel biological mechanisms that improve functional tendon regeneration is therefore an important unmet need.

Toward that end, we previously established a neonatal mouse model of functional tendon regeneration after Achilles tendon transection without repair (7). We found that neonatal tendon regeneration was characterized by early proliferation of tendon cells, followed by their recruitment into the injury site to form a tenogenic neo-tendon. In the absence of tendon cell recruitment, functional properties were not restored (8). Using a similar injury model in adult mice, we also found that adult tendon healing was characterized by fibrotic scar formation, minimal tendon cell proliferation, and persistently reduced functional properties (7), consistent with studies from other groups (9–11).

Although the immune response is a key aspect of tendon healing, the specific immune cells and signals underlying tendon healing have yet to be fully defined. It is generally accepted that tendon healing is initiated by infiltration of pro-inflammatory macrophages (M1-like macrophages) followed by resolution of inflammation by alternatively activated macrophages (M2-like macrophages) (12–15). However, the role of these macrophage populations in promoting or inhibiting tendon healing is still unclear (16). While therapeutic strategies that enhance the presence of M2-like macrophages appear to promote functional tendon healing (17, 18), other studies suggest that increased M2-like macrophage polarization can also drive fibrotic scar formation (19). Most of these studies are correlative however and there are still very few studies that directly test the role of macrophages in tendon healing via ablation or depletion. One well-established method for macrophage depletion is delivery of liposomal clodronate (20). In adult mouse models of Achilles tendon transection and repair, macrophage depletion by clodronate resulted in reduced cell proliferation and improved mechanical properties (21, 22). Similarly, clodronate depletion in an ACL reconstruction rat model showed improved mechanical properties at the tendon-bone interface (23). Despite these results suggesting that macrophages impair adult functional healing, there is also evidence indicating that increased macrophage recruitment can also improve tensile properties after adult tendon injury (24). To date, the role of neonatal macrophages in tendon healing have not been investigated.

To address these open questions, we determined the immune response during neonatal tendon healing by screening immune-related gene expression and tested the requirement for macrophages using an established genetic model of macrophage depletion to inducibly ablate macrophages. We hypothesized that neonatal tendons would exhibit an anti-inflammatory immune response after injury and that macrophage depletion would impair functional regeneration.

## MATERIALS AND METHODS

### Experimental procedures

For experiments, the transgenic Macrophage-Fas-Induced Apoptosis (MaFIA) mouse line (25) and ScxGFP tendon reporter line (26) were used. Although the MaFIA line allows GFP-detection of *Csfr1*-expressing monocytes and macrophages, we were only able detect MaFIA^GFP^ by flow cytometry and not by fluorescence microscopy of either spleen or tendon. Flow cytometry was therefore used to quantify macrophage depletion in mice containing only the MaFIA allele. For other experiments, ScxGFP was incorporated into the MaFIA background to enable ScxGFP cell visualization with macrophage depletion (MaFIA/ScxGFP). EdU was given at 0.05 mg 2 hours prior to harvest to label proliferating cells. Global macrophage depletion was carried out using the homodimerizer AP20187 at 10 mg/kg in carrier solution by subcutaneous injection (Cat. # 635058, Clontech) with carrier-treated controls (4% EtOH, 10% PEG-400, 1.7% Tween20) injected in parallel. Littermates from the same litter were evenly split between AP20187-treatment or carrier-treatment. Full Achilles tendon transection without repair was carried out in neonates at P5, with male and female mice distributed evenly between groups. All procedures were approved by the Institutional Animal Care and Use Committee at Mount Sinai.

### Flow cytometry quantification

Single cell suspensions were generated from Achilles tendons of MaFIA mice by enzymatic digestion with a solution containing 1 mg/mL collagenase I (Cat. # LS004196, Worthington Biochemical) and 5 mg/mL collagenase IV (Cat. # LS004188, Worthington Biochemical) for 2.5 hrs at 37C. Cells were then stained 1:1000 with DAPI (Cat. # D1306, ThermoFisher) in 2% FBS in PBS to detect live cells. Flow cytometry was carried out on an LSRIIA machine (BD Sciences) using FACSDiva software and analyses using FCS Express 7. Gating for MaFIA^GFP^ was performed using control cells isolated in parallel from a wild type mouse with no GFP.

### Immunofluorescence, EdU detection, and fluorescence microscopy

Limbs were immediately fixed in 4% PFA after harvest for 24 hours at 4°C and then decalcified in 50 mM EDTA for 1-2 weeks at 4°C. To embed, limbs were incubated in 5% sucrose (1 hour) and 30% sucrose (overnight) at 4°C and then embedded in OCT medium (Cat. # 23-730, Fisher Scientific). Transverse cryosections (12 μm) were collected in alternating slides. Immunostaining for macrophages was carried out using an antibody against the global macrophage marker F4/80 (Cat. # 14-4801, Affymetrix) and secondary detection by Cy5 (Cat. # 712-175-150, Jackson ImmunoResearch). EdU labeling was detected with the Click it EdU kit in accordance with manufacturer’s instructions (Cat. # C10340, Life Technologies). Fluorescence images of EdU and ScxGFP were acquired using the Zeiss Axio Imager with optical sectioning by Apotome. Cell quantification was performed in Image J software using the CellCounter plugin. All images for quantifications were taken at the same exposure and image manipulations applied equally across samples.

### RNA isolation, reverse transcription, and qRT-PCR

RNA was extracted from Achilles tendons using Trizol/chloroform. cDNA was synthesized by reverse transcription using the SuperScript VILO master mix (Cat. # 11755050, Invitrogen). qPCR was carried out by Mount Sinai’s core facilities using SYBR PCR Master Mix (Cat. # 4309155, Thermo Fisher) and gene expression calculated using the 2^−ΔΔCt^ method relative to *Gapdh* and carrier-treated control tendons at D3. Primers for *Il1β* (FWD: AGTTGACGGACCCCAAAAGAT; REV: GTTGATGTGCTGCTGCGAGA) and *Il10* (FWD: ATTTGAATTCCCTGGGTGAGAAG; REV: CACAGGGGAGAAATCGATGACA) were used. Tendon gene primers were previously described (7).

### Gait analysis

On D28, mice were gaited at 10 cm/s for 3-4 s using the DigiGait Imaging System (Mouse Specifics Inc.). A high-speed digital camera was used to capture forelimb paw positions and parameters previously validated for mouse Achilles tendon injury (Swing, Brake, and Propel) were then calculated (7). All parameters were normalized to Stride length to account for differences in animal size since male and female mice were used.

### Biomechanical testing

For biomechanical testing, Achilles tendons were dissected at D56, wrapped in PBS-soaked gauze and frozen at −20C until time of testing. Tensile testing was performed in PBS at room temperature using custom 3D printed grips to secure the calcaneus bone and Achilles tendon (27, 28). Tendons were preloaded to 0.05N for ~1 min followed by ramp to failure at 1%/s.

### Statistical analysis

Quantitative results are presented as mean±standard deviation. For tendon macrophage flow cytometry, functional properties, and gene expression, two way ANOVA was used for comparisons with independent variables of treatment (carrier vs AP20187) or injury; where significance was detected, posthoc testing was then carried out (Graphpad Prism). Flow cytometry of spleen macrophages and all other quantitative analyses were analyzed using Students t-tests. Significant outliers were detected using Grubb’s test (Graphpad Prism). Significance was determined at p<0.05.

## RESULTS

### Neonatal tendon injury activates an early pro-inflammatory immune response

To define the immune response associated with neonatal tendon regeneration, we carried out a gene expression screen using the Taqman Array Mouse Immune Response panel comprising 96 genes. Neonatal injured and contralateral uninjured Achilles tendons were screened at 3 days post-injury (D3), prior to recruitment of tenogenic cells in the gap space. While we initially expected neonatal injured tendons to exhibit a minimal immune response, we found 27 genes were upregulated after injury (≥2-fold, p<0.05, **Table 1**) with 4 genes were downregulated (≤-2-fold, p<0.05, **Table 2**). Upregulated markers included genes associated with macrophages and monocyte/macrophage chemotaxis (*Ccl2, Ccl3, Ccr2, Ccr7*, Cd68) and T-cells (*Tbx21, Cd3e, Ctla4, Cxcr3*). Surprisingly, there appeared to be a robust inflammatory response as we detected upregulation of several pro-inflammatory cytokines (*Il6, Il1β, Il12β, Il18*); however, at this timepoint the anti-inflammatory cytokine *Il10* was also upregulated. In addition to upregulation of *Tgfβ1*, we detected significant downregulation of *Ski*, which is a known negative regulator of TGFβ signaling. Collectively, our qPCR screen indicates that neonates mount a robust pro-inflammatory immune response after injury that may be driven by recruitment of macrophages and T-cells.

**Table 1:** Differentially expressed genes detected by Taqman Array Mouse Immune Response panel in response to neonatal tendon injury at D3.

| **Gene** | **Fold Change (Up-regulated)** | **p value** |
| --- | --- | --- |
| *Gzmb* | 183.2675 | 0.045003 |
| *Ccl5* | 38.95 | 0.000215 |
| *Il6* | 23.5654 | 0.026617 |
| *Prf1* | 23.4663 | 0.006136 |
| *Tbx21* | 22.367 | 0.003557 |
| *Sele* | 18.1162 | 0.028513 |
| *Tnfrsf18* | 14.0241 | 0.009632 |
| *Cd3e* | 13.907 | 0.006046 |
| *Ctla4* | 13.7629 | 0.00171 |
| *Vcam1* | 13.5854 | 0.013771 |
| *Il1β* | 11.77 | 0.000642 |
| *Il10* | 11.3756 | 0.002664 |
| *Ccl3* | 10.1302 | 0.017634 |
| *Ccl2* | 10.0745 | 0.024852 |
| *Ccr7* | 9.366 | 0.000269 |
| *FasI* | 7.1758 | 0.001069 |
| *Il12β* | 6.952 | 0.02003 |
| *H2-Eb1* | 6.9387 | 0.000538 |
| *Cd80* | 5.7704 | 0.030404 |
| *Ccr2* | 5.7292 | 0.005281 |
| *Cxcr3* | 4.0058 | 0.010695 |
| *Csf2* | 3.737 | 0.009016 |
| *Stat1* | 3.5504 | 0.040912 |
| *Il18* | 3.2386 | 0.045934 |
| *Cd68* | 3.1542 | 0.003324 |
| *Hmox1* | 2.4951 | 0.011452 |
| *Tgfb1* | 2.1161 | 0.011681 |
| **Genes** | **Fold Change (Down-regulated)** | **p value** |
| *Nos2* | -2.3753 | 0.005565 |
| *Ski* | -2.23 | 0.000631 |
| *Vegfa* | -2.1495 | 0.009757 |
| *Ace* | -2.0735 | 0.040496 |

### Increased macrophages and macrophage localization with tenocytes after neonatal tendon injury

Analysis of the pan-macrophage marker F4/80 in transverse cryosections from uninjured tendon (P8 post-birth) showed tendon-resident macrophages are normally sparsely detected in the tendon periphery as well as in fascia surrounding the tendon (**Figure 1A, B**). By D3 post-injury, we observed a dramatic increase in F4/80+ cells surrounding the injured tendon. Comparison of sections taken from the tendon stub far from the cut site as well as sections close to the cut site showed that F4/80+ macrophages were only located in close proximity to ScxGFP+ tendon cells in sections near the injury site (**Figure 1C**).

**Figure 1:**
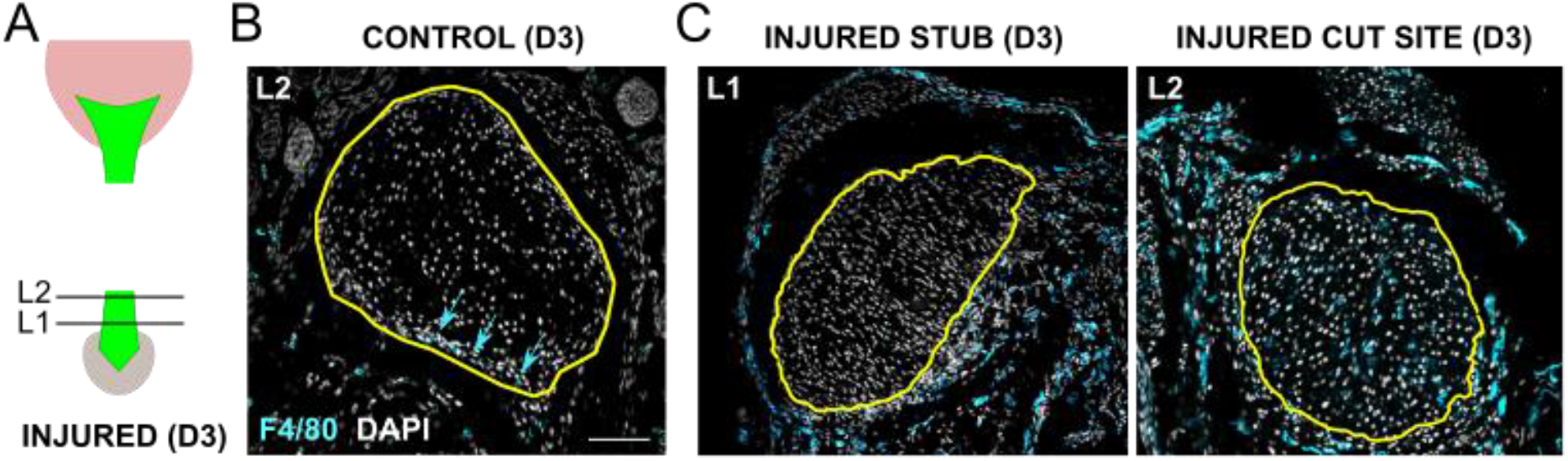
Macrophages are recruited after neonatal tendon injury and localize to cut site tendon cells. (**A**) Transverse cryosections were collected from tendon stub (L1) and cut site (L2) levels shown in schematic. Immunostaining for pan-macrophage marker F4/80 of (**B**) uninjured control and (**C**) injured tendon at D3 post-injury. Yellow outline indicates tendon region. Blue arrows point to sparse F4/80+ tendon-resident macrophages residing in tendon periphery. Scale bar: 100 μm.

### Repeated depletion of macrophages during postnatal growth resulted in adverse systemic effects unrelated to tendon healing

The close proximity of macrophages to tenocytes suggested potential interactions between these cells. We therefore tested the functional role of macrophages on neonatal tendon regeneration using the Macrophage Fas-Induced Apoptosis transgenic mutant (MaFIA) in which monocytes and macrophages express a mutant human FK506 binding protein. Macrophages could therefore be inducibly and reversibly depleted by delivery of the homodimerizer AP20187. For all experiments, AP20187 or carrier was injected for three consecutive days (P2, P3, P4) prior to Achilles tendon transection at P5 to deplete monocytes and macrophages prior to injury. We then tested three different regimens of macrophage depletion after P5 injury, including twice a week, once a week, or a single booster dose at d7 post-injury. When AP20187 was delivered twice a week following injury, we observed swelling of both injured and uninjured hindlimbs of treated mice and impaired growth (**Figure 2A-C**). Once a week treatment improved survival of mice out to nearly d56, however restricted growth was observed beginning at d35 (**Figure 2D-F**). Swelling was also observed in both injured and uninjured hindlimbs of surviving mice at d56. Finally, we treated mice with a single booster injection at d7 and observed no difference in growth and no swelling of hindlimbs (**Figure 3A-C**). Since the single booster treatment showed minimal systemic effects and has also been reported in other studies (29), we carried out subsequent analyses using this treatment regimen. Analysis of ablation efficiency by flow cytometry of MaFIA^GFP+^ showed 86% reduction in macrophages at d3 in the injured tendon and 93% reduction in the spleen following AP20187 treatment (**Figure 3D**).

**Figure 2:**
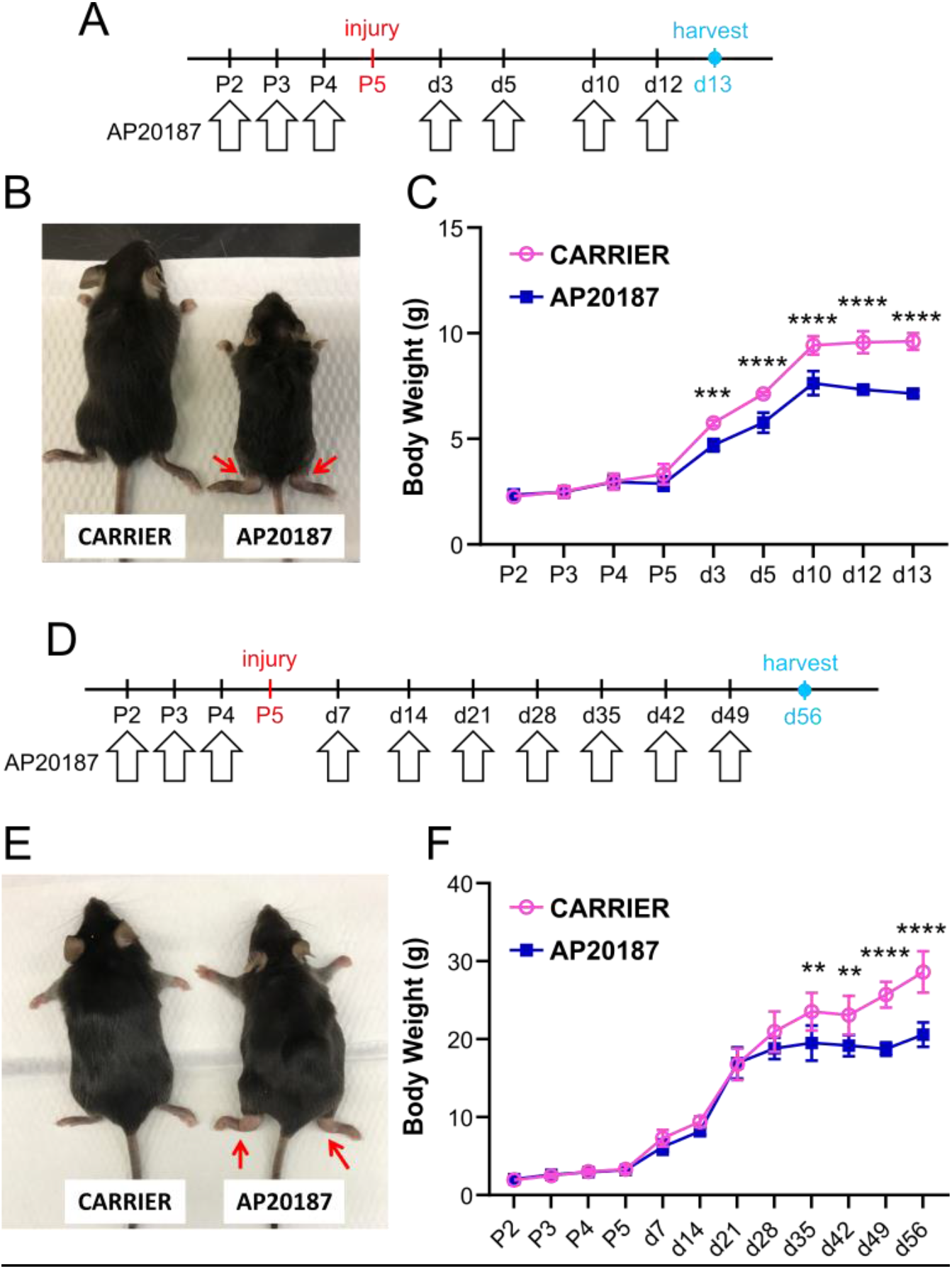
Prolonged and repeated depletion of macrophages results in adverse systemic effects. (**A**) Schematic showing twice a week dosing regimen of AP20187 after P5 Achilles tendon injury. (**B**) AP20187-treated mice were smaller and showed abnormal swelling of ankles at D13 (red arrows) with (**C**) impaired weight gain (n=3-5). (**D**) Schematic showing once a week dosing regimen of AP20187 after P5 Achilles tendon injury. (**E**) AP20187-treated mice also developed abnormal swelling of ankles at D56 (red arrows) and (**F**) impaired weight gain at later timepoints (n=3-5). ** p<0.01; *** p<0.001 **** p<0.0001.

**Figure 3:**
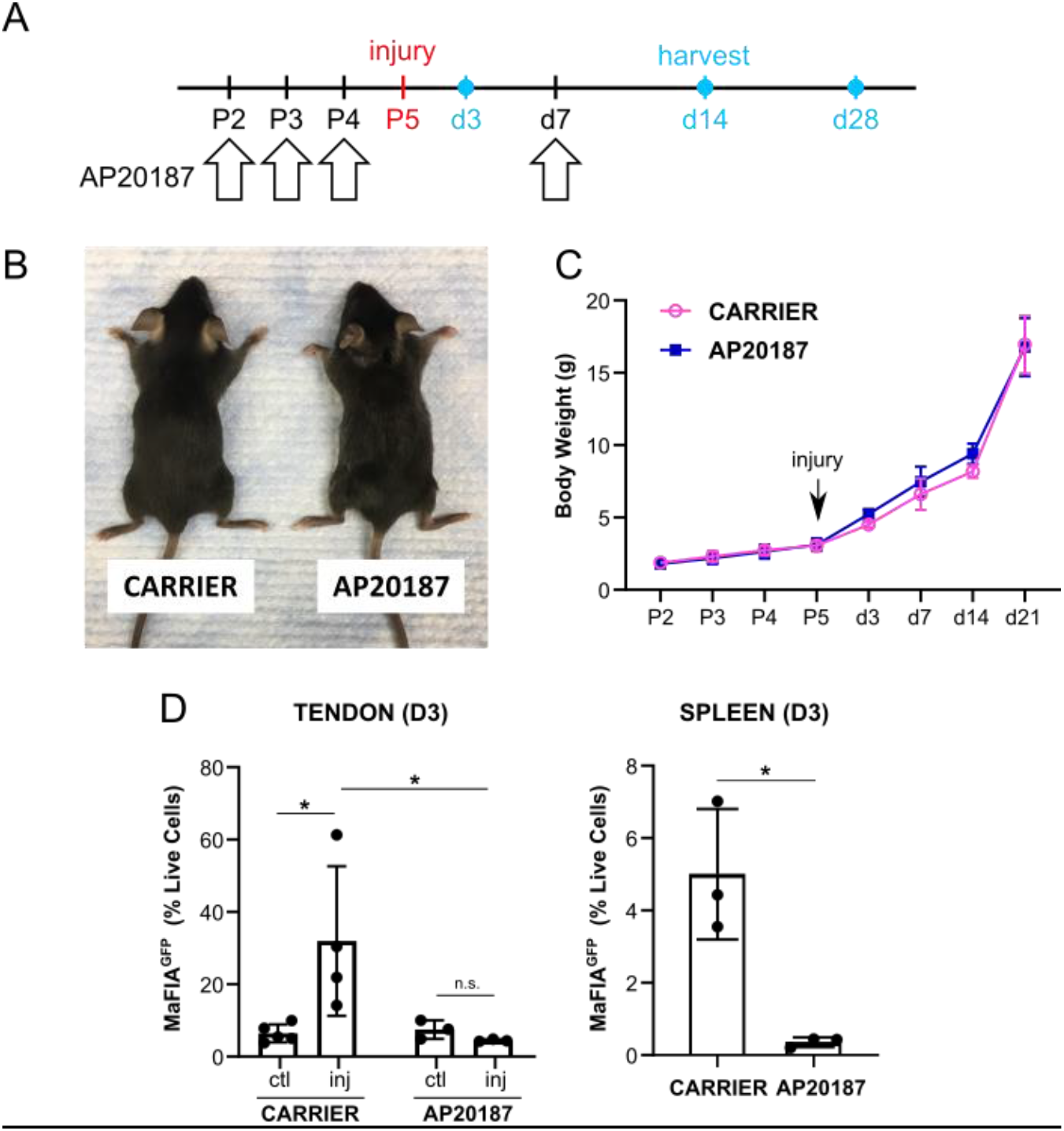
Validation of macrophage depletion by AP20187. (**A**) Schematic showing a single booster dose of AP20187 at D7 after P5 Achilles tendon injury. (**B**) AP20187-treated mice did not demonstrate swelling of hindlimbs compared to carrier-treated at D28. (**C**) Weight gain was comparable at all timepoints (n=4-9). (**D**) Flow cytometry quantification of MaFIA^GFP^ cells at D3 in tendon and spleen. * p<0.05.

### Tendon mechanical properties are not restored after macrophage depletion

To determine functional healing when macrophages were ablated, we carried out gait analysis and tensile testing. In previous studies, we identified swing, brake, and propel parameters (normalized to stride length) were affected by injury (7, 8). Analysis of gait at 28 showed no differences with AP20187-treatment, either compared to contralateral uninjured control or compared to carrier treatment for any of the three parameters (**Figure 4**). However, direct tensile testing of tendons at D56 showed impaired functional restoration with macrophage ablation after injury. Stiffness was reduced in AP20187-treated injured tendons compared to the contralateral control tendon and max force was reduced compared to both contralateral control and carrier injured tendons (**Figure 5**). There was no difference in mechanical properties between control and injured limbs with carrier treatment. These results indicate that macrophages are required for functional neonatal tendon healing.

**Figure 4:**
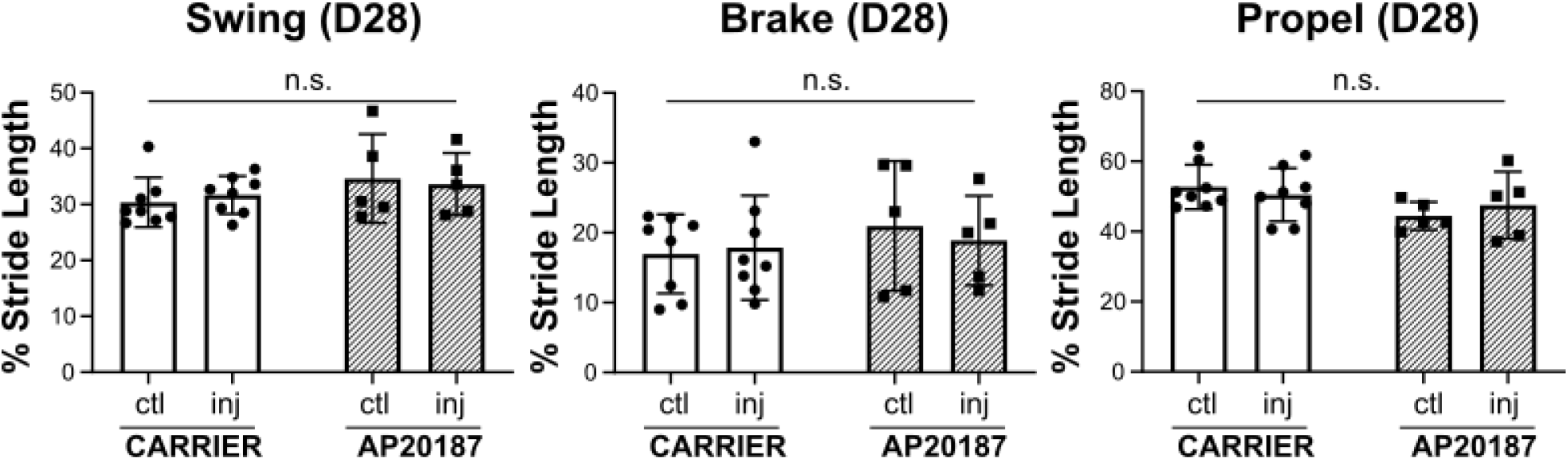
Recovery of gait after tendon injury was not affected by macrophage depletion. Gait analysis showed functional recovery of all gait parameters (Swing, Brake, and Propel) at D28 post-injury regardless of treatment. n.s. indicates p>0.05. n=5-8.

**Figure 5:**
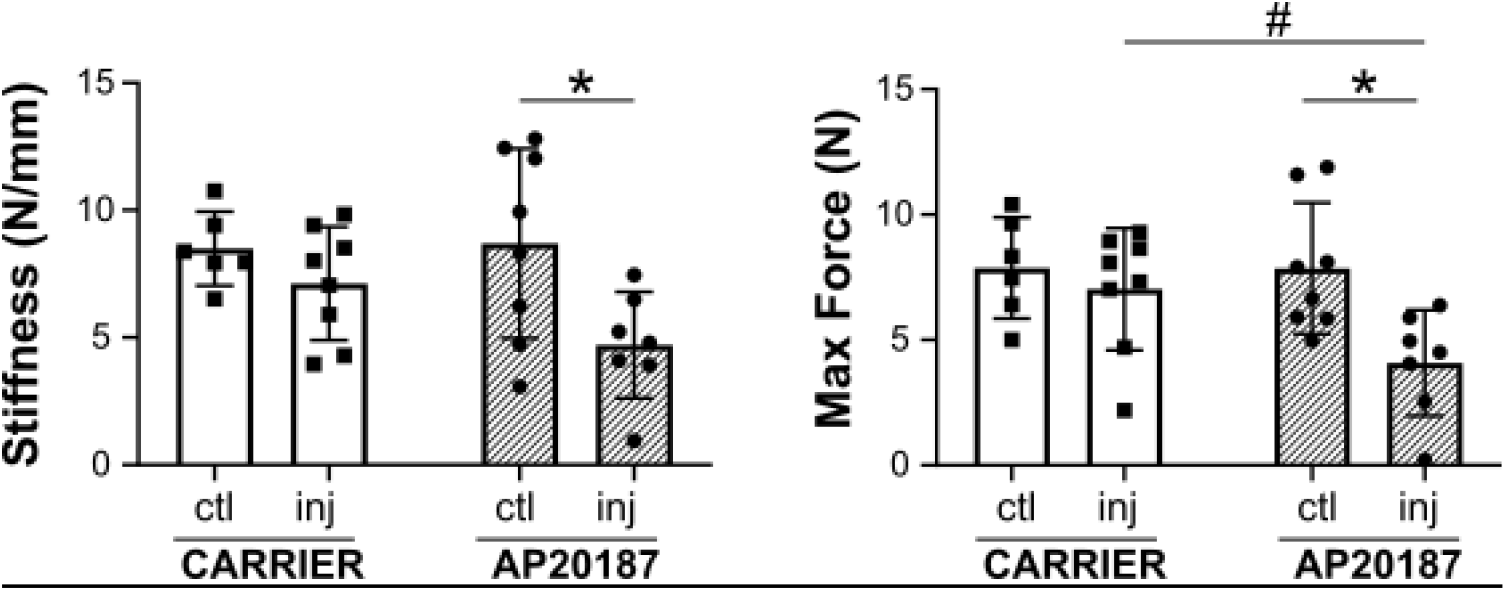
Mechanical properties are impaired after tendon injury with macrophage depletion. Tensile testing of Achilles tendons at D56 post-injury showed AP20187-treated tendons did not fully restore stiffness and max force after injury compared to contralateral controls (n=6-8). * p<0.05 # p<0.1.

### Early ScxGFP+ cell proliferation and subsequent ScxGFP+ neo-tendon formation is impaired with macrophage depletion

To understand the cellular deficits that might be associated with functional loss, we analyzed proliferation at D3 by EdU incorporation (2 hrs) and ScxGFP neo-tendon formation at D14 (**Figure 6A**). With carrier treatment, intense proliferation of both ScxGFP+ and ScxGFP-cells was observed at D3 in the tendon cut site (**Figure 6B**). In contrast, minimal proliferation occurred with AP20187 treatment and the number of proliferating ScxGFP+ cells was reduced. By D14, a robust ScxGFP+ neo-tendon was formed in carrier-treated mice, however macrophage depletion resulted in significantly reduced ScxGFP+ area (**Figure 6C**). Taken together, this suggests that macrophage depletion resulted in defects in tendon cell proliferation leading to reduced ScxGFP cell recruitment and neo-tendon formation.

**Figure 6:**
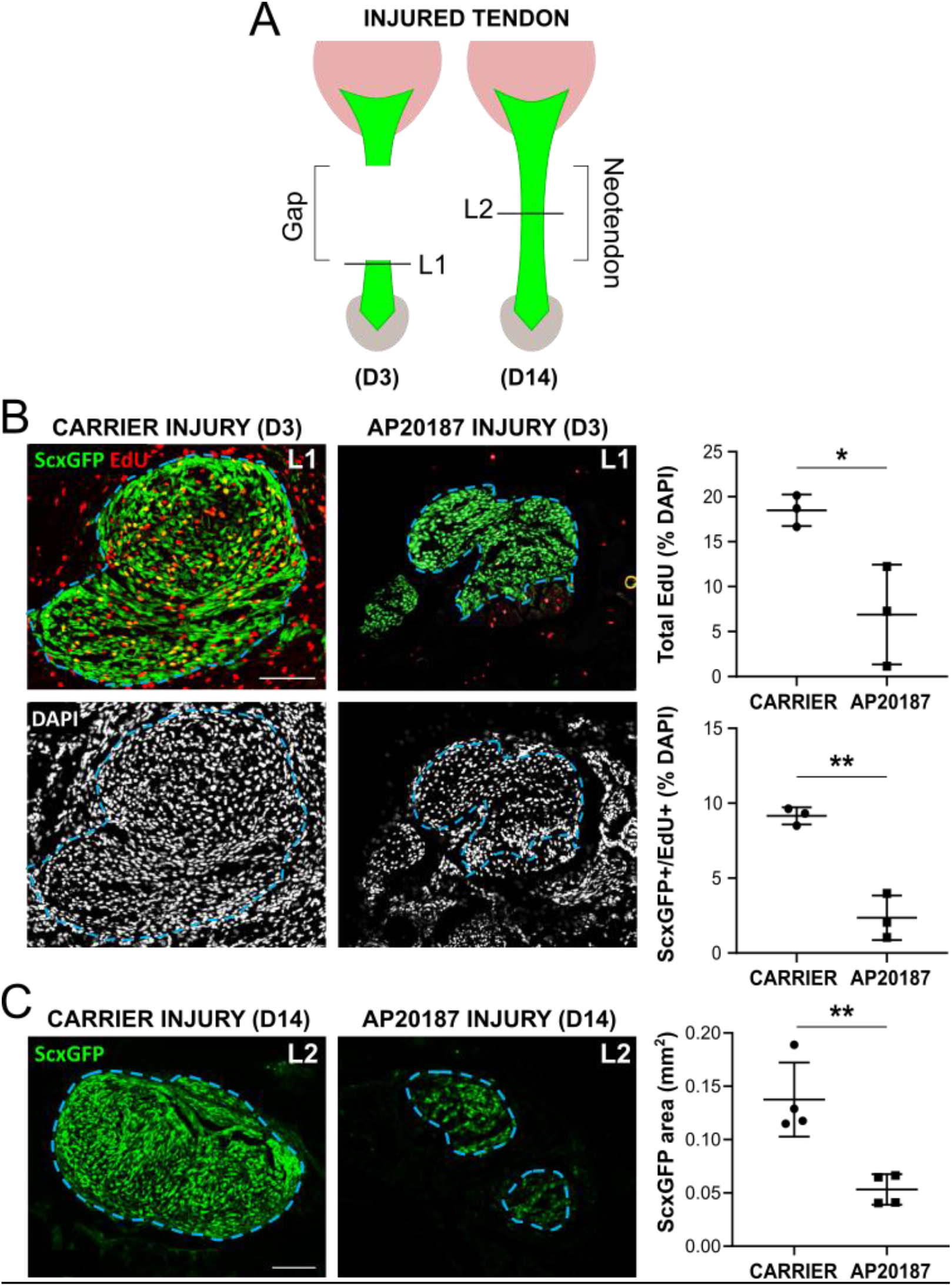
Reduced proliferation and ScxGFP neo-tendon formation after tendon injury with macrophage depletion. (A) Transverse cryosections were collected from tendon cut site at D3 (L1) and neo-tendon region at D14 (L2). (B) EdU and ScxGFP imaging and cell quantification at D3 post-injury (n=3). (C) ScxGFP area quantification at D14 post-injury (n=4). Blue dashed outlines show tendon region of interest based on ScxGFP expression. * p<0.05 ** p<0.01.

### Macrophage depletion results in altered gene expression of the contralateral control tendon

Although the neo-tendon area was reduced with macrophage depletion, we still observed a population of ScxGFP+ cells recruited at D14, indicating a positive tendon phenotype. To better define the tenogenic phenotype of these cells, we carried out gene expression analysis at D3 and D14 by qPCR. Expression of known markers *Scx*, *Tnmd*, and *Mkx* was reduced in carrier treated injured limbs relative to contralateral controls at D3 (**Figure 7A**), consistent with our prior studies (8). In AP20187-treated limbs, injury induced a less predictable response. While *Tnmd* expression decreased with injury, *Scx* and *Mkx* levels were not significantly different (p>0.1). At D14, a small reduction in *Tnmd* and *Mkx* expression was still observed in carrier-treated injured limb relative to control with no difference in AP20187-treated limbs (**Figure 7B**). However, this was due to a decrease in *Tnmd* and *Mkx* in AP20187-treated control limbs (compared to carrier-treated controls). Unexpectedly, *Scx* expression in AP20187-treated control limbs was decreased at D3 relative to carrier-treated control limbs, but dramatically increased by D14 relative to all other groups (**Figure 7A, B**). This may indicate a potential rebounding effect, although ScxGFP expression was not visibly altered (not shown). There was no difference in tendon gene expression between carrier-treated and AP-treated injured limbs at any timepoint.

**Figure 7:**
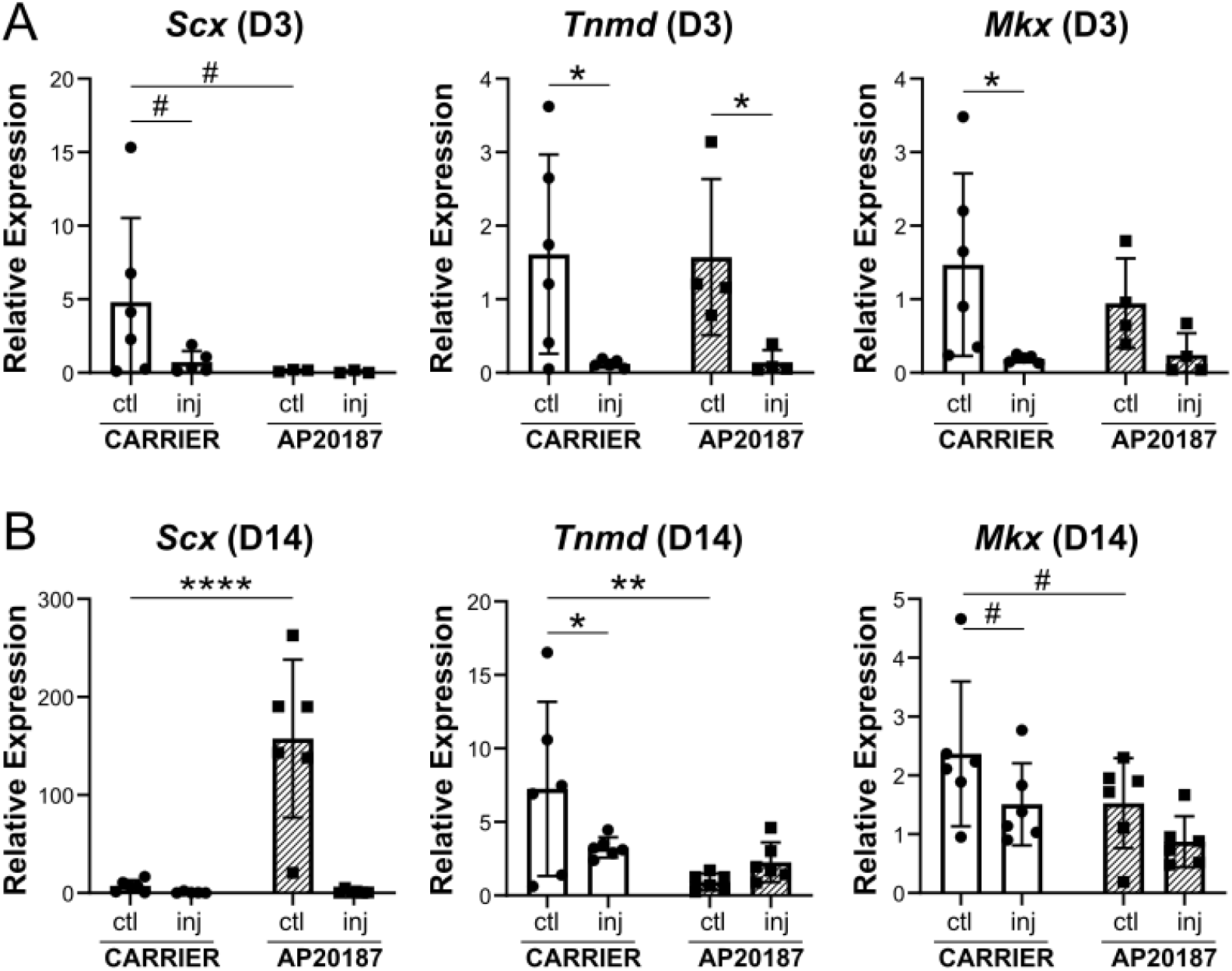
Tendon gene expression is altered with macrophage depletion. Expression of tendon genes Scx, Tnmd, and Mkx at (**A**) D3 and (**B**) D14 post-injury (n=4-6). * p<0.05 ** p<0.01 **** p<0.0001 # p<0.1.

### Macrophage depletion may enhance early inflammation

To determine the effects of macrophage depletion on local inflammation, *Il1β* and *Il10* expression was determined by qPCR at D3 and D14. At D3, *Il1β* expression was increased with injury compared to controls, regardless of treatment (**Figure 8A**). Levels of *Il1β* was higher in AP20187-treated injured limbs relative to carrier-limbs, although the difference did not reach significance (p=0.08) due to high variability. At D14, there was persistent *Il1β* upregulation in carrier-treated injured limbs, but no difference with AP20187-treatment (**Figure 8B**). There was no difference in *Il10* expression at any timepoint for either treatment or injury. Overall, this data suggests that there may be somewhat early elevation of inflammation with macrophage depletion.

**Figure 8:**
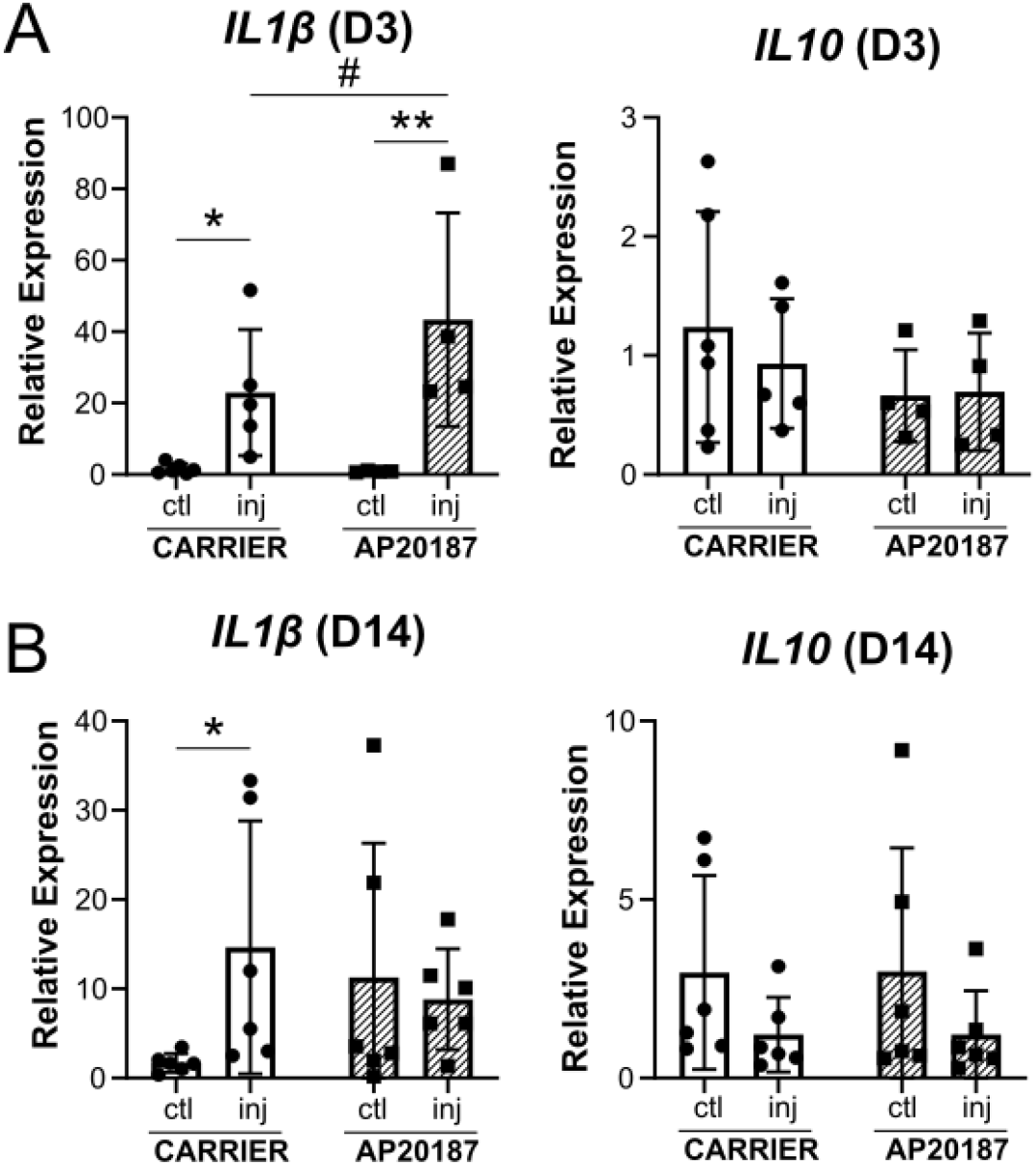
Expression of cytokines with macrophage depletion. Expression of *Il1β* and *Il10* at (**A**) D3 and (**B**) D14 post-injury (n=4-6). * p<0.05 ** p<0.01 # p<0.1.

## DISCUSSION

In this study, we screened for immune gene expression markers after tendon injury and found upregulation of several pro-inflammatory markers, suggesting that early inflammation is a key feature of neonatal tendon regeneration. This was contrary to our initial hypothesis, as it was previously suggested that scar-less fetal regeneration occurred in a permissive immune environment characterized by minimal inflammation (30–32). Increased inflammation in the fetal environment resulted in failed regeneration of fetal skin and tendons (30, 33); this could be rescued by dampening inflammation via IL-10 over-expression (30). Interestingly, there is also evidence that adult tissues are unable to regenerate when placed in the fetal environment, suggesting that a permissive immune environment may be necessary but not sufficient to fully transform adult scar-mediated healing (34). Although fetal tendons transplanted into the adult environment were still able to regenerate (35), immunodeficient hosts were used in these studies and the fetal tendons therefore did not experience a normal adult immune response to injury. To date, there are still few studies focused on the immune regulation of neonatal regeneration. However, it is well appreciated that the neonatal immune system is less mature compared to adult and is biased toward type 2 immunity in response to infections (36, 37). Many categories of neonatal immune cells (including neutrophils, natural killer cells, monocytes, and others) also show substantial functional differences and numbers compared to adults (37), reflecting the unique immune environment of the neonate. While we observed enhanced expression of pro-inflammatory markers, analyses were limited to a single early timepoint. Ongoing studies will define the temporal dynamics of inflammation and its resolution (or persistence) in neonates in direct comparison with adult counterparts after tendon injury. It may be that neonatal inflammation is reduced compared to adults or is more rapidly resolved.

Depletion of macrophages during early healing resulted in impaired functional properties likely due to reduced ScxGFP tendon cell proliferation, recruitment, and neo-tendon formation. The close proximity of macrophages to tendon cells at D3 suggests that secreted factors from macrophages may stimulate tendon cell proliferation. While TGFβ is an attractive candidate based on our gene expression array screen and has been implicated in other macrophage depletion models of tendon healing (21, 23) as well as fetal tendon healing (35, 38), our previous studies inhibiting TGFβ signaling showed no effect on tendon cell proliferation at D3 (8). The molecular regulators for early cell proliferation is therefore still unknown. These factors may be identified by transcriptional profiling by RNA sequencing of isolated MaFIA^GFP^ macrophages or by single cell RNA sequencing.

One major limitation of this study includes the timing of AP20187 delivery and macrophage depletion. The goal of this study was to determine broadly the role of macrophages; we therefore depleted macrophages prior to injury with an additional booster injection at D7. While previous research in adult tendons suggest that M2-like macrophages do not emerge until D28 in the healing timeline (12), it is possible we also depleted M2-like macrophages. In our study, we observed reduced ScxGFP+ cell proliferation at D3, during the time when M1-like macrophages are expected to be dominant. This suggests that M1-like macrophages are important for stimulating tendon cell proliferation. While other studies also showed a requirement for macrophages in cell proliferation (21, 22), this is the first study to distinguish tendon cells (ScxGFP+) from exogenous or dedifferentiated (ScxGFP−) cells. Our finding that inflammation may be crucial for the early healing response is consistent with other research showing that inhibition of inflammation too early in tendon healing can have detrimental outcomes on functional properties (39). Future studies will determine the temporal dynamics and functions of distinct macrophage sub-populations recruited during neonatal tendon healing and test depletion strategies targeting early (D0-D3), middle (D5-D9), and late (D20-24) phases. Temporal depletion of macrophages will also determine whether impaired neo-tendon formation at D14 is due solely to reduced proliferation of tendon cells or whether there is an additional contribution of tendon cell migration.

Although carrier-treated mice restored gait and mechanical function after injury, gene expression analyses indicated a potential alteration or delay in healing at early timepoints. Our previous studies generally showed restored or upregulation of tendon markers by D14, however carrier-treated injured tendons in this study maintained decreased expression of *Tnmd* and *Mkx*. This may be due to an effect of the carrier solution on tendon healing. Similarly, Il1β was also elevated in carrier-treated injured tendons at D14; in unpublished data from the lab using untreated mice, we generally observe resolution of neonatal inflammation at this timepoint (not shown). Interestingly, there was also an effect of AP20187-treatment on the contralateral uninjured tendon, in terms of tendon gene expression. While flow cytometry did not show a difference in macrophage number in control tendons with AP20187-treatment, we cannot rule out the possibility that there may be some degree of tendon-resident macrophage apoptosis leading to low-level local inflammation independent of injury. We also observed nearly complete macrophage depletion in the spleen, which may alter systemic circulating factors that may affect the contralateral control limb. In future studies, the impact of local vs systemic effects will be distinguished by local injection of AP20187 directly to the injured or control limb.

Finally, although macrophages are key players in the healing process, they are not the only immune cells that may regulate regeneration. During muscle regeneration, specific T cell subpopulations (Th1, Th2 and Tregs) have been shown to induce or impede regenerative healing (40–43). Our qPCR array screen also identified markers associated with T cells, consistent with previous studies showing their recruitment during tendon healing and in human tendinopathy (44–46). Ongoing work will test the role of T cells in tendon regeneration and scar formation.

## ACKNOWLEDGMENTS

This work was supported by NIH/NIAMS (R01AR069537, R56AR076984) to AHH and F31AR073626 to DK. We gratefully thank Dr. Ronen Schweitzer for providing ScxGFP mice and the qPCR Core at Mount Sinai for their assistance with this project.

## AUTHOR CONTRIBUTIONS

KH, DK, and AHH contributed to research conception and experimental design. KH, DK, AM, KY, and PN contributed to data collection. Data analyses were performed by KH, DK, and AHH. Manuscript was written by AHH and all authors contributed to edits and approve this submission.

